# Diversity analysis of amp gene sequences in the ‘*Candidatus* Phytoplasma meliae’

**DOI:** 10.1101/2020.06.01.128413

**Authors:** Franco D. Fernández, Luis R. Conci

## Abstract

Phytoplasmas are plant pathogenic bacteria transmitted by insects. As endosymbiotic bacteria that lack a cell wall, their membrane proteins are in direct contact with host cytoplasm. In phytoplasmas the immunodominant membrane proteins (IDPs), are the most abundant proteins of the cell membrane. The antigenic membrane protein (Amp), one of the three types of IDPs, is characterized by a positive selection pressure acting in their extracellular domain. In South America, the ‘*Candidatus* Phytoplasma meliae’ has been associated to chinaberry yellows disease. In the present work, we describe for the first time the structure, phylogeny and selection pressure of amp gene in sixteen ‘*Candidatus* Phytoplasma meliae’ isolates. Our results indicate that amp gene sequences preserve the structure, large extracellular domain flanked by to hydrophobic domains in the N- (signal peptide) and C-termini (transmembrane), previously described in its orthologues and high divergence in the amino acids residues from extracellular domain. Moreover, a positive selection pressure was detected predominantly in this region confirming previous reports.

## Introduction

Phytoplasmas are cell wall-less bacteria that inhabit sieve cells in the phloem tissue of infected plants and are transmitted from plant-to-plant by phloem-feeding insect vectors, principally leafhoppers (Zhao et al. 2015). These pathogens are associated with plant diseases in several hundred plant species, including many important food, vegetable and fruit crops, ornamental plants, timber and shade trees (Bertaccini and Lee, 2019). In South America, China berry trees (*Melia azedarach* L) are affected by two different phytoplasmas, ‘*Candidatus* Phytoplasma meliae’ (group 16SrXIII, subgroups –C and –G) (Fernández et al. 2016) (Figure 1A) and ‘*Candidatus* Phytoplasma pruni’ (group 16SrIII, subgroup B) (Galdeano et al. 2004). In Argentina, ‘*Ca*. P. meliae’ is restricted to North-East region while ‘*Ca*. P. pruni’ has a wider distribution covering different regions of the country (Arneodo et al. 2007; Fernandez 2015). The differential distribution of these two phytoplasmas could be linked to the distribution of its insect vector, considering that each species of phytoplasmas establishes a unique relationship with the insect that has the ability to transmit it. Nevertheless, in this case this hypothesis has not been confirmed yet. In this context, the study of membrane proteins is a reliable approach to understand the molecular dialogue between insect vectors and phytoplasmas. This group of pathogens lacks a cell wall, thus their membrane proteins are in direct contact with the host cytoplasm (Konnerth et al. 2016). The immunodominant membrane proteins (IDPs) are a group of proteins that comprises a major portion of total cellular membrane proteins in phytoplasmas (Kakizawa et al. 2004). To date, three non-orthologous IDPs types have been described: Imp (immunodominant membrane protein), Amp (antigenic membrane protein) and IdpA (immunodominant membrane protein A) (Kakizawa et al. 2006a). The Amp protein is constituted by a large extracellular domain flanked by two hydrophobic domains in the N- (signal peptide) and C-termini (transmembrane) (Arashida et al. 2008; Barbara et al. 2002; Kakizawa et al. 2006a). Previous studies have shown great variability in the extracellular domain accompanied by high selection positive pressure (Fabre et al. 2011; Kakizawa, et al. 2006b). This selection pressure is suggested to be associated with the key role that it plays in the interaction of phytoplasmas with insect vectors (Suzuki et al. 2006). So far, studies carried out with the Amp protein have been only described in aster yellows group phytoplasmas (16SrI, ‘*Ca*. Phytoplasma asteris’) and Stolbur (16SrXII, ‘*Ca*. Phytoplasma solani’). In South America, there are no reports about the Amp protein in ‘*Ca*. Phytoplasma meliae’ and related phytoplasmas of the 16SrXIII group (Mexican periwinkle virescence). In this scenario, the goal of this work was to describe the main features of Amp in diverse geographical isolates of ‘*Ca*. Phytoplasma meliae’ present in Argentina to study its variability and selection pressure processes.

**Figure 1.**
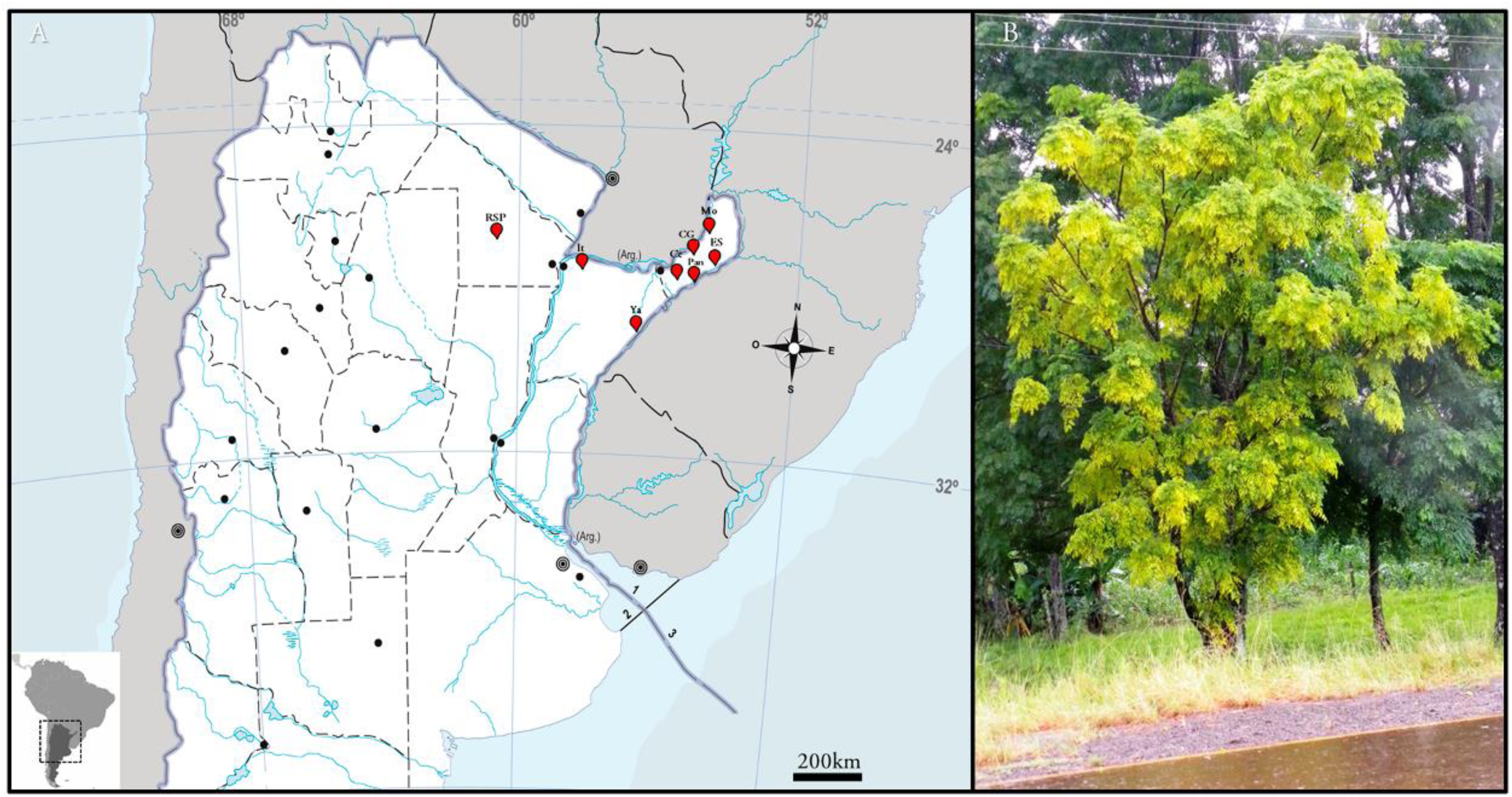
‘*Candidatus* Phytoplasma meliae’ affecting China berry trees in Argentina. A: Partial map of Argentina showing the sampling points (red), Campo Grande (CG), El Soberbio (ES), Montecarlo (Mo) and Panambí (Pan) from Misiones province, Itatí (It) and Yapeyú (Ya) from Corrientes province, and Roque Saenz Peña (RSP) from Chaco province; B: China berry tree showing typical symptoms of yellowing.

## Materials and methods

### Sample source

Total DNA from sixteen (n=16) chinaberry tree naturally infected with ‘*Candidatus* Phytoplasma meliae’ were used in molecular analyzes. This DNA collection was obtained from different geographical locations situated in the northeast of Argentina (Table 1, Figure 1A). For DNA extraction CTAB protocol (Doyle and Doyle 1990) was used. Detection and identification of ‘*Candidatus* Phytoplasma meliae’ was accessed by PCR and PCR-RFLP as described previously (Fernández et al. 2016). Briefly, PCR detection was conducted using universal primers P1/P7 (Deng and Hiruki 1991) and R16F2n/R16R2 (Lee et al. 1994) in direct and nested reactions. Nested PCR amplicons (1.2kb) were subjected to digestion using *Mse*I, *Hpa*II, *Rsa*I and *Hae*III (NEB, USA) endonucleases and RFLP profiles were compared to reference patterns of subgroups 16SXIII-G and 16SrIII-B.

**Table 1:** ‘Ca. Phytoplasma meliae’ samples used in this work. * Latitude/Longitude (decimal)

| *Ca. meliae* isolate | Location | Province | Coordinates* | #accession |
| --- | --- | --- | --- | --- |
| ChTY-25.1 | Campo Grande (CG) | Misiones | -27.207945°, -54.979693° | MG905031.1 |
| ChTY-Ce3 | Cerro Azul (Ce) | Misiones | -27.633535°, -55.497152° | MG905032.1 |
| ChTY-27.1 | El Soberbio (ES) | Misiones | -27.29549°, -54.196343° | MG905019.1 |
| ChTY-27.6 | El Soberbio (ES) | Misiones | -27.29549°, -54.196343° | MG905020.1 |
| ChTY-27.9 | El Soberbio (ES) | Misiones | -27.29549°, -54.196343° | MG905030.1 |
| ChTY-Mo3<sup>R</sup> | Monte Carlo (Mo) | Misiones | -26.566667°, -54.783333° | MG905024.1 |
| ChTY-Mo2 | Monte Carlo (Mo) | Misiones | -26.566667°, -54.783333° | MG905026.1 |
| ChTY-Mo6 | Monte Carlo (Mo) | Misiones | -26.566667°, -54.783333° | MG905021.1 |
| ChTY-30.1 | Panambí (Pan) | Misiones | -27.7223°, -54.914895° | MG905029.1 |
| ChTY-IT26 | Itatí (It) | Corrientes | -27.266667°, -58.25° | MG905022.1 |
| ChTY-IT27 | Itatí (It) | Corrientes | -27.266667°, -58.25° | MG905027.1 |
| ChTY-IT25 | Itatí (It) | Corrientes | -27.266667°, -58.25° | MN699857.1 |
| ChTY-Ya4 | Yapeyú (Ya) | Corrientes | -29.469611°, -56.817444° | MN699858.1 |
| ChTY-RS3 | Roque Saenz Peña (RSP) | Chaco | -26.783333°, -60.45° | MG905023.1 |
| ChTY-RS12 | Roque Saenz Peña (RSP) | Chaco | -26.783333°, -60.45° | MG905025.1 |
| ChTY-RS13 | Roque Saenz Peña (RSP) | Chaco | -26.783333°, -60.45° | MG905028.1 |

### Amplification and sequencing of groEL-amp-nadE region

DNA from ‘*Ca*. Phytoplasma meliae’ (isolate ChTYXIII-Mo3) was used as reference DNA for sequencing genomic fragment containing *groEL* (partial)-*amp* (complete)-*nadE* (partial) genes. Firstly, a degenerate primer pair (*groEL-Fw1/nadE-Rv2*) (Table 2) was designed manually based in the sequences of *groEL* (cpn60) and *nadE* genes from related ‘*Ca*. Phytoplasma species’ available in GenBank. Amplification of 3.2 kb was obtained and directly sequenced from both ends using the same primers. Based on these sequences new specific primers pair (*groEL-ChTYFw1/nadE-ChTYRv1*), which amplified a putative fragment of 2.0 kb, were designed using Primer3 implemented in Geneious R.10 (Biomatters, USA). PCR amplifications were conducted in a final volume of 50 µl, containing 1.5U of Dream® Taq polimerase (Fermentas, Lituania), 0.4 µM of each primer, 100 µM of dNTPs and 1X buffer Dream Taq (2 mM MgCl2). For 2.0 kb amplification PCR conditions used were, 3 minutes 94°C for initial denaturation and 35 cycles of 94°C/1minute, 58°C/1 minute and 72°C/3 minutes, with final extension of 72°C for 10 minutes. The PCR product (2.0 kb) was purified using S-400 HR columns (GE, UK) and cloned in pGEM T-Easy system (Promega, USA) according to the manufacturer instructions. The complete sequence of 2.0 kb amplicon was obtained by *primer walking* strategy in three different clones (Macrogen, Korea).

**Table 2:**
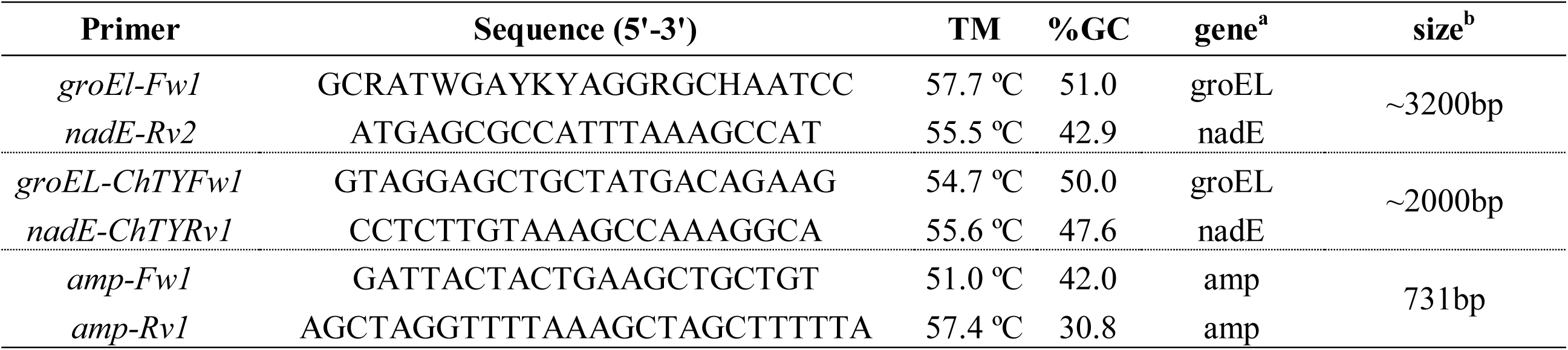
List of primer used in this work. ^a^ target gene, ^b^ PCR fragment size

### Structural analysis

Open reading frames were estimated using ORF Finder in Geneious R.10 software. Annotation of amino acidic deduced sequences was performed using BLASTp (nr, BLOSUM62, word size 6). For Amp-ORF signal peptide sequence and the cleavage site were predicted with the program SignalIP v5.0 (http://www.cbs.dtu.dk/services/SignalP/) as well as the presence of the transmembrane domains with TMHMM v2.0 program (http://www.cbs.dtu.dk/services/TMHMM/). Also, the conserve domains were analyzed by CD-Search online tool (https://www.ncbi.nlm.nih.gov/Structure/cdd/wrpsb.cgi).

### Phylogeny

For phylogeny reconstruction multiple alignments of amino acid sequences were conducted using MAFFT (L-INS-i, 200PAM/K=2, gap open penalty=1.53, offset value=0.123) (Katoh and Toh 2008), from Amp sequences obtained in this work and from related phytoplasmas groups (16SrI and 16SrXII) available from GenBank. The evolutionary history was inferred by using the Maximum Likelihood method based on the Le Gascuel model. Multiple alignments of 16S rDNA gene sequences were performed using MUSCLE (window size=5, gap open score= −1) and evolutionary history was inferred using the Maximum Likelihood method based on the General Time Reversible model. In both cases, bootstrap (1,000 repetitions) was performed for statistical support. Initial tree(s) for the heuristic search were obtained automatically by applying Neighbor-Join and BioNJ algorithms to a matrix of pairwise distances estimated using a JTT model, and then selecting the topology with superior log likelihood value. All evolutionary analysis were conducted in MEGA7 (Kumar et al. 2016).

### Selection pressure on *amp* gene

In order to elucidate the selection pressure acting in the *amp* gene, fifteen new ‘*Ca*. Phytoplasma meliae’ isolates were sequenced (Table 1). A new set of primers, *amp*Fw1-*amp*Rv1 were designed based on *groEL-amp-nadE* (2.0 kb) sequence previously described (ChTYXIII-Mo3). These primers amplified a putative fragment of ∼0.7 kb containing the entire sequence of the *amp* gene. Cloning and sequencing were conducted as previously described. For each ‘*Ca*. Phytoplasma meliae’ isolate 3 different clones were bidirectionally sequenced and consensus sequences (3X minimum coverage) were obtained using Geneious R10 and deposited in GenBank. For the target gene (*amp*) the synonymous (dS) and non-synonymous (dN) nucleotide substitution rates were calculated. The dN/dS ratios and the null hypothesis of no selection (H0: dN=dS) versus the positive selection hypothesis (H1: dN>dS) were calculated using Nei–Gojobori method in a codon-based Z selection test implemented in MEGA7 software (Kumar et al. 2016). The variance of the difference was computed using the bootstrap method (1,000 replicates). In case of positive selection dN/dS ratio must be >1 and p-value for the Z-test < 0.05 (Masatoshi and Sudhir 2000). Maximum Likelihood computations of dN and dS were also conducted using HyPhy software package (Pond et al., 2005). The statistic test dN - dS is used to detect codons that have undergone positive selection, where dS is the number of synonymous substitutions per site (s/S) and dN is the number of nonsynonymous substitutions per site (n/N). A positive value for the statistic test indicates an overabundance of nonsynonymous substitutions. Normalized dN - dS for the statistic test is obtained using the total number of substitutions in the tree (measured in expected substitutions per site) which were also calculated in order to compare different data sets. Tajima’s test of neutrality (Tajima 1989) was also conducted using MEGA7. Three set of data were used in this work, ‘*Ca*. Phytoplasma meliae’ *amp* sequences data set (n=16) (this paper); ‘*Ca*. Phytoplasma solani’ *STAMP* sequences data set (n=15) (Fabre et al. 2011) and ‘*Ca*. Phytoplasma asteris’ *amp* sequences data set (n=13) (Kakizawa et al. 2006a).

## Results

### Amplification and sequencing of groEL-*amp*-nadE fragment

Using the primer pair *groEL-Fw1/nadE-Rv2* a ∼2 kb PCR fragment was amplified in all ‘*Ca*. Phytoplasma meliae’ samples (16/16). No amplification product was obtained from healthy chinaberry samples (data not show). PCR amplification of ‘*Ca*. Phytoplasma meliae’ isolate ChTYXIII-Mo3 (reference strain) (Fernández et al. 2016) was selected for sequencing. A final consensus sequence of 1,975 bp was obtained and deposited in GenBank under accession MG905024. ORF estimation revealed that the sequenced fragment contains two incomplete ORFs (ORF-1_1-630_ and ORF-3_1694-1975_) and one complete (ORF-2_750-1226_) (Figure 2.A). BLASTp analysis showed that ORF-1 encodes a protein homologue to Chaperonine GroEL (*groEL*) (86.89% identity, E=1e-125, accession: CBL82429.1) and the ORF-3 encodes a NAD synthetase (*nadE*) (76.09% identity, E=2e-45, accession: BAG16386.1). The protein encoded by ORF-2 (474bp-158aa) showed an identity of 41% (E=2e-24) with Amp of ‘*Candidatus* Phytoplasma japonicum’ (BAG16385). In the intergenic region 1 (IG1_631-749_) putative transcription signals (−35: TTTATG; −10: TAATAGGTT) were found while in the intergenic region 2 (IG2_1227-1693_) a putative transcription terminator (TGTTTTTAAAAAGCTAGCTTTAAAACCTAGCTTTTTTTCTTTATTC) was also found. Comparison of *groEL-amp-nadE* genomic fragment showed a high conservation in ORFs corresponding to groEL and nadE proteins, while for amp gene, lower identity values were observed mainly in the central region (Figure 2.A). These results confirmed the synthetic organization of the genes flanking amp in the order 5’-*groEL-amp-nadE*-3’ as previously reported in others ‘*Ca*. Phytoplasmas species’ (Arashida et al. 2008; Fabre et al. 2011; Kakizawa et al. 2006b) or STOLBUR phytoplasmas (Fabre et al., 2011).

**Figure 2.**
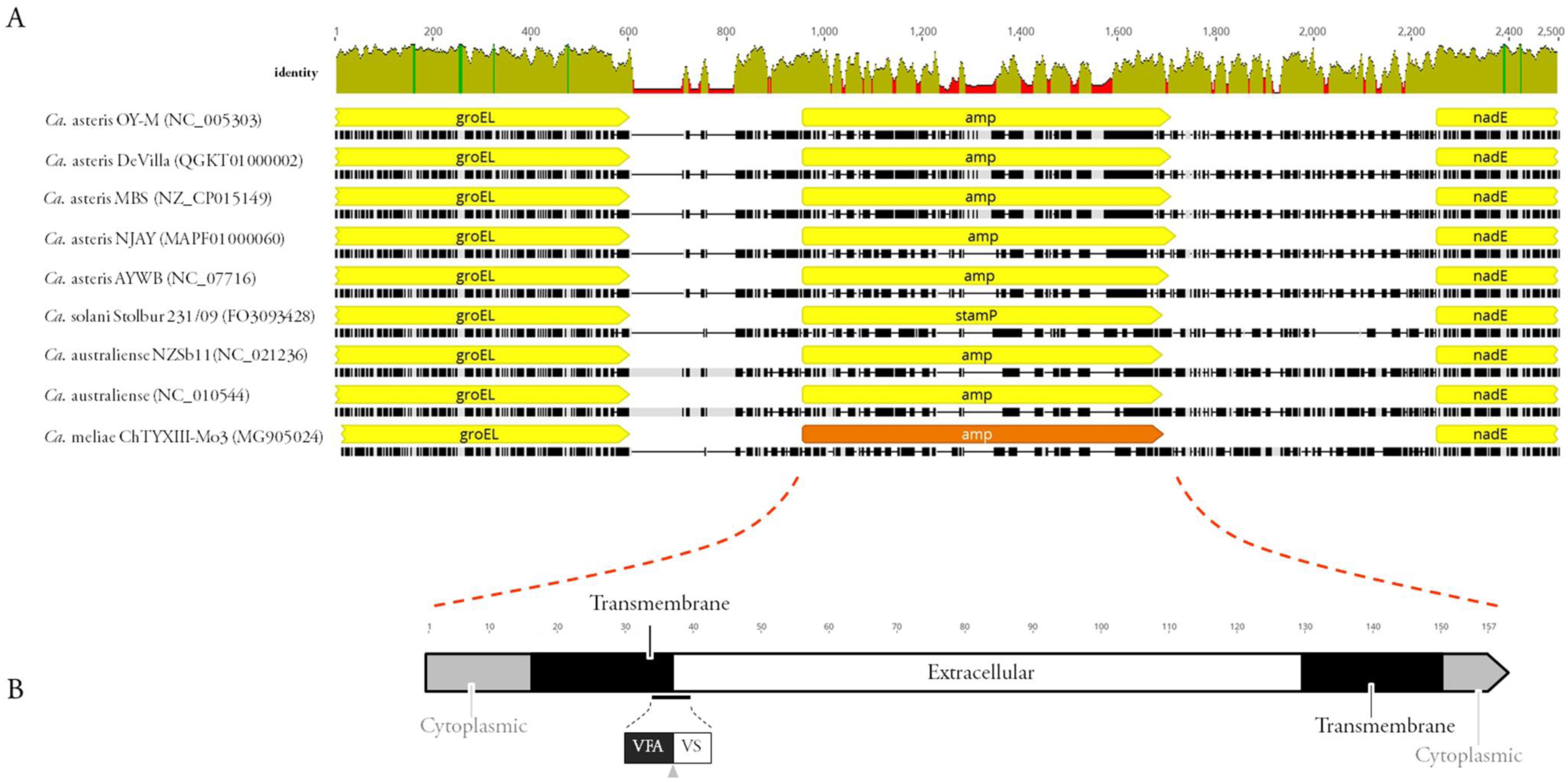
Genetic context of *Ca*. Phytoplasma meliae’Amp of ‘. A: multiple alignments of groEL-amp-nadE loci in related ‘*Ca*. Phytoplasma species’. Identity values are showed. B: structure of putative Amp protein, cytoplasmic domains (grey), transmembrane domains (black), extracellular domain (white), putative cleavage motif (amino acid residues VFA-VS) (black line).

### Structural analysis of amp protein

The deduced amino acid sequence for the Amp was 158 aa, with an estimated molecular weight of 17.07 kDa. Regarding its composition, the Amp is rich in alanine residues (12%), serine (12%), valine (9.13%) and lysine (14.6%). Based on SignalP-5.0 analysis, residues corresponding to the signal peptide (1-35) (Signal peptide (Sec/SPI), Likelihood = 0.9587), and a cleavage site of the putative protein between residues 35-36 (VFA-VS/Probability = 0.8813) (Figure 2.B) were identified. Two transmembrane regions, at the N-termini (signal peptide) (residues 13-35) and C-termini region (residues 131-152) (Figure 2.B), and an extracellular domain (residues 36-130) were also identified. pSORTb prediction located Amp as cytoplasmic membrane protein (score = 9.87). Phyto-Amp conserved domain (Cdd: pfam15438) was also recorded in the interval 1-103 (E= 0.02) supporting that this protein is an orthologous of previously described antigenic membrane protein (Amp). Despite the conserved structural organization of Amp domains (TM-extracellular-TM) among orthologues described in different *‘Ca*. Phytoplasma species’, we observed that some ‘*Ca*. Phytoplasma asteris’ isolates presented a somewhat larger extracellular region (positions 77-116 and 132-172) (Figure 3.B).

**Figure 3.**
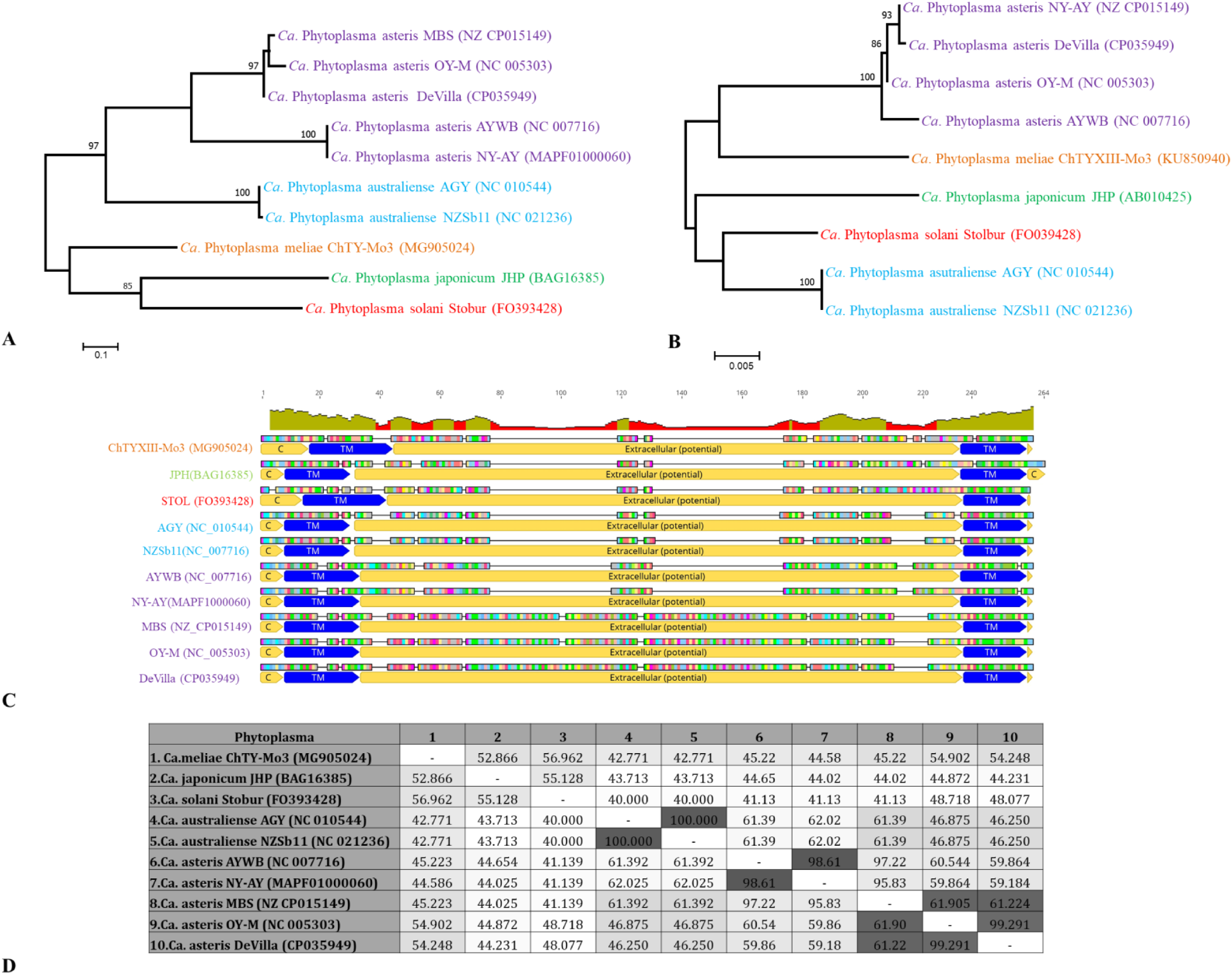
Comparative analysis of Amp. A-B: Phylogenetic relationships inferred from analysis of Amp and 16Sr RNA gene sequence, respectively, using the Maximum Likelihood method implemented in MEGA 7. The bootstrap consensus tree inferred from 1,000 replicates is taken to represent the evolutionary history of the taxa analyzed. Sequences obtained in this work are in bold. The scale bar represents the number of nucleotide substitutions per site. Bootstrap values > 70% are shown in the nodes. ‘*Ca*. Phytoplasma species’ are written in different colors. C: Multiple alignment of Amp protein sequence from diverse ‘*Ca*. Phytoplasma species’, different domains in each sequence are illustrated with yellow (extracellular and cytoplasmic-C) or blue (transmembrane) colors. D: amino acids identity values expressed as % and heatmap.

### Phylogeny

Phylogenetic analyses were performed from both Amp and 16S rDNA sequences. For Amp, ML tree shows that ‘*Ca*. Phytoplasma meliae’ grouped in the same clade with ‘*Ca*. Phytoplasma solani’ and ‘*Ca*. Phytoplasma japonicum’ (Figure 3.A). Meanwhile, the general topology of 16S rDNA ML-tree does not consistently correspond to that described for Amp, since the groupings generated do not share the same-clustered taxonomic groups (Figure 3.B), and ‘*Ca*. Phytoplasma meliae’ was grouped more closely with different species of ‘*Ca*. Phytoplasma asteris’.

### Selection pressure on *amp* gene

Sixteen (n=16) ‘*Candidatus* Phytoplasma meliae’ *amp* gene sequences were used in selection pressure analysis. Multiple alignments of 474 positions (158 codons) were evaluated and dN-dS calculated for each codon (Supplementary material, Table S1). Fourteen codons showed values of dN-dS > 0, which would indicate that they are under a positive selection pressure (overall dN-dS=2.884, p=0.005), of them, nine were found to encode amino acids in the extracellular region (Table 3, Figure S1). The same analysis was also performed with two sets of data from population studies conducted with Amp in other phytoplasmas species. The first data set (consisting of 15 sequences) corresponded to the immunodominant protein STAMP (Fabre et al. 2011) characterized in various European isolates of the STOLBUR phytoplasmas. The other data set (consisting of 13 sequences) corresponded to the Amp characterized in various isolates of ‘*Ca*. asteris’ related phytoplasmas (Kakizawa et al. 2006a). In both cases, the dN-dS values were calculated, and their position within the sequence (Transmembrane domains, Extracellular domains). For STAMP protein, out of 154 codons analyzed, 19 had values of dN-dS > 0 (overall= 2.226, p=0.028). Sixteen out these nineteen codons were located in the extracellular region (Table 3). On the other hand, the Amp-asteris protein, presented 62 codons, over a total of 225, with dN-dS values > 0 (overall=4.764, p< 0.001). Within these 62 codons, 57 were associated to extracellular domain (Table 3).The results of these analyses determined that the highest number of codons with dN-dS values > 0 occurred in the extracellular region (Figure 4), which would indicate a positive selection pressure acting on this domain.

**Table 3:**
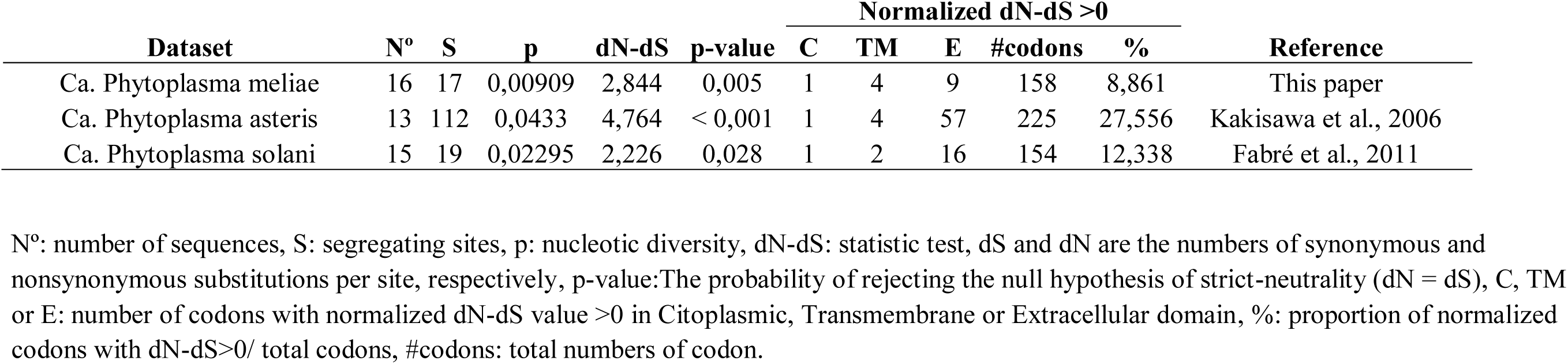
Selection pressure analysis in AMP proteins from three different ‘*Ca*. Phytoplasma specie’ data sets.

| Dataset | N° | S | p | dN-dS | p-value | Normalized dN-dS >0 |  |  |  |  | Reference |
| --- | --- | --- | --- | --- | --- | --- | --- | --- | --- | --- | --- |
|  |  |  |  |  |  | C | TM | E | #codons | % |  |
| Ca. Phytoplasma meliae | 16 | 17 | 0,00909 | 2,844 | 0,005 | 1 | 4 | 9 | 158 | 8,861 | This paper |
| Ca. Phytoplasma asteris | 13 | 112 | 0,0433 | 4,764 | < 0,001 | 1 | 4 | 57 | 225 | 27,556 | Kakisawa et al., 2006 |
| Ca. Phytoplasma solani | 15 | 19 | 0,02295 | 2,226 | 0,028 | 1 | 2 | 16 | 154 | 12,338 | Fabré et al., 2011 |

N°: number of sequences, S: segregating sites, p: nucleotic diversity, dN-dS: statistic test, dS and dN are the numbers of synonymous and nonsynonymous substitutions per site, respectively, p-value: The probability of rejecting the null hypothesis of strict-neutrality ( $dN = dS$ ), C, TM or E: number of codons with normalized dN-dS value >0 in Citoplasmic, Transmembrane or Extracellular domain, %: proportion of normalized codons with $dN-dS > 0$ / total codons, #codons: total numbers of codon.

**Figure 4.**
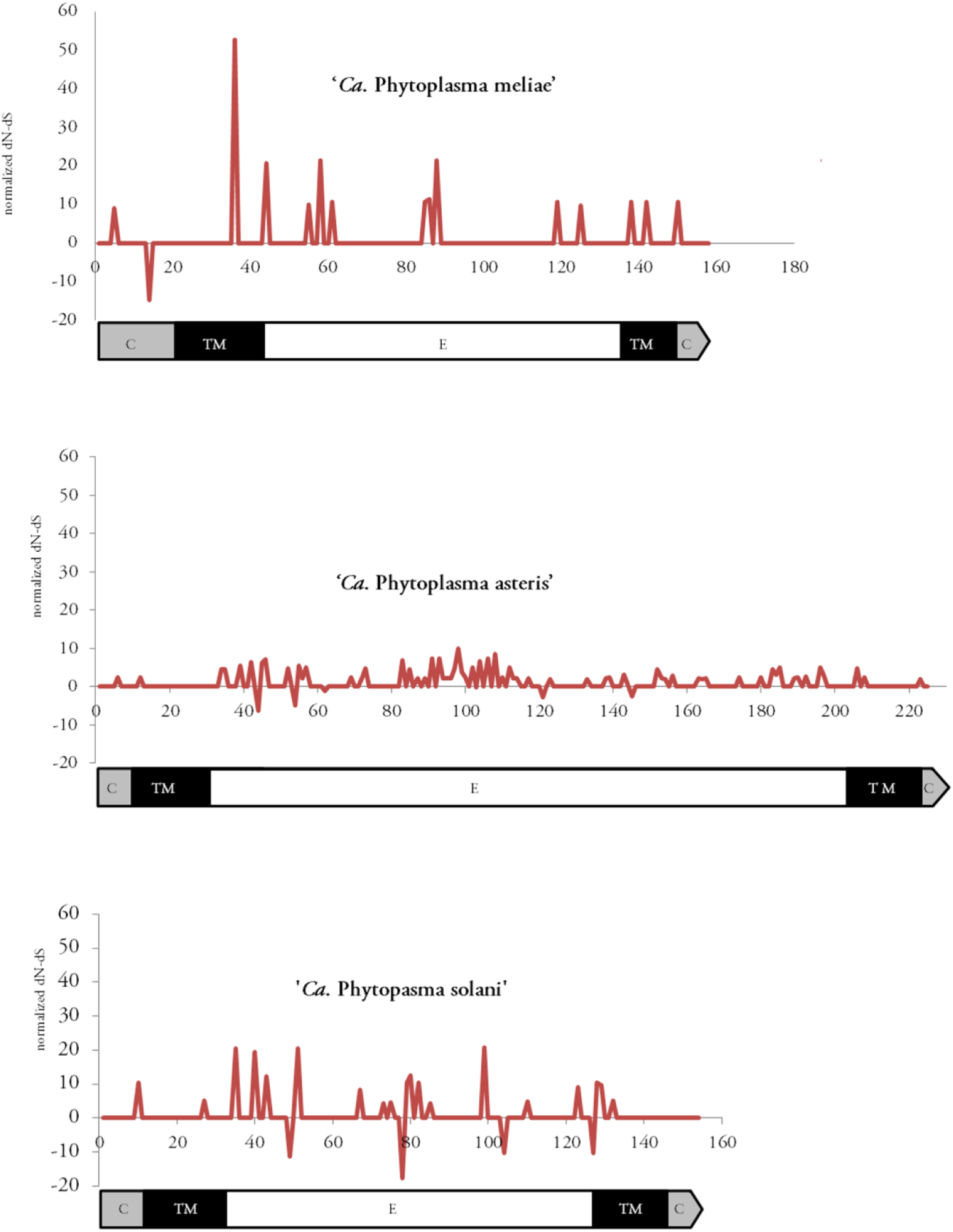
Selection pressure acting on Amp protein. The results are displayed for three data sets, amp-ChTY (16 sequences), STamp-STOLBUR (9 sequences) and amp-asteris (14 sequences). In each data set the position of the codon is plotted on the X axis, and on the Y axis the standardized dN-dS value corresponding to each one of those codons. Domains of Amp protein are also illustrated at scale in the X axis.

## Discussion

In this work we described and characterized for the first time amp gene in sixteen isolates from ‘*Ca.* Phytoplasma meliae’ derived from different geographical regions in Argentina. Previous studies have shown the high conservation in *groEL-amp-nadE* operon from diverse ‘*Ca.* Phytoplasma species’ (Andersen et al. 2013; Arashida et al. 2008; Barbara et al. 2002; Coetzee et al. 2019; Fabre et al. 2011; Kakizawa et al. 2006b; Sparks et al. 2018). Our analysis showed that this 5’-*groEL-amp-nadE*-3’ locus organization is also conserved in ‘*Ca.* Phytoplasma meliae’. The general structure of Amp consisted in a large extracellular hydrophilic domain flanked by two hydrophobic domains in the N- and C-termini. While the C-terminal domain contained a transmembrane region, which could serve as an anchor to phytoplasma cellular membrane, the N-terminal domain included a signal peptide which is probably cleaved during protein maturation (Arashida et al. 2008). Amp described in the present work consisted in 160-158 aa, with a molecular weight of ∼17 kDa, two hydrophobic domains located in the N-terminal (signal peptide) and C-terminal ends (transmembrane), and a signal peptide cleavage (VFA-VS) within residues 33-37. A central hydrophilic region was also inferred as the mayor portion of Amp-meliae protein. These features indicated that the characterized protein is an orthologous of Amp described in other phytoplasmas species. Comparative analysis with others “C. Phytoplasma” showed that Amp of ‘*Candidatus* Phytoplasma meliae’ had a low aa homology (23.33%-37.74% identity), with the central hydrophilic region being the most variable. This is consistent with what has been previously described in the aster group phytoplasma (Kakizawa et al. 2006a) and the Stolbur group (Fabre et al. 2011). It has also been reported that the Amp protein is also divergent in size (Arashida et al. 2008) and its extracellular region may vary between ∼175 aa (*Ca.* asteris strains MBS, DeVilla and OY-M) to ∼100 aa (*Ca.* solani, *Ca.* australiense and *Ca.* asteris strains AYWB and NYAY). In the case of Amp from *Ca.* meliae, the extracellular region is composed of 94 aa, which is more closely linked to those phytoplasmas that have the smallest size in that domain. Likewise, reconstruction of the phylogeny of *Ca.* meliae Amp protein has also linked it more closely with the phytoplasmas *Ca.* solani and *Ca*. japonicum than *Ca.* australiense and *Ca.* asteris species. This association was not consistent with those obtained for the highly conserved 16S rDNA gene, indicating the presence of selective pressure acting on the Amp. The impact of positive selection on the rate of protein evolution is evident in only a small fraction of proteins, mainly those subjected to recurrent positive selection that is typically associated to host–pathogen interactions (Zhang et al. 2015). Several studies strongly suggest that positive selection is acting on IDPs (Amp, Imp and idpA) (revised in Konnerth et al. 2016). In Amp, most of amino acids subjected to positive selection pressure are located in the extracellular domain (Fabre et al. 2011; Kakizawa et al. 2006a). The results obtained in this work have allowed us to confirm this pattern, since Amp in ‘*Ca*. Phytoplasma meliae’ appears to be subjected to a positive selection pressure (overall dN-dS> 0) resulting in a diversifying positive selection exerting in this portion on the gene. The strong selection pressure and high divergence described for Amp and other IDPs proteins, suggest that they might be playing a key role in the phytoplasma-host molecular interaction (Kakizawa et al. 2009; Kakizawa et al. 2006a). In fact, it has been shown that the Amp in OY-M phytoplasma forms a complex with actin microfilaments in leafhoppers which determine insect-vector specificity (Suzuki et al. 2006). It has also been shown that Amp of CYP phytoplasma specifically binds to α and β subunits of ATP synthases of insect vectors (Galetto et al. 2011). The role of this protein was also evaluated with pre-feeding assays of two CYP vector with specific Amp-antibody which resulted in significant decrease in the acquisition efficiency (Rashidi et al. 2015). Moreover, the Amp is somehow involved in the specific crossing of the gut epithelium, as well as salivary gland colonization, during the early phases of vector infection with CYP (Pacifico et al. 2015). Blocking IDPs protein using a specific scFv (Le Gall et al. 1998) or antibody (Pacifico et al. 2015) in plants or an insect vector was highly effective in reducing phytoplasma infection in both hosts. Recently a RNAi strategy was implemented via microinjection of muscle actin and ATP synthase β dsRNAs in adult insects of *E. variegatus* which caused an exponential reduction in the expression of both genes and also a significant decrease in survival rates (Abbà et al. 2019). Considering the aforementioned characteristics, the Amp protein constitutes an interesting target not only for the development of resistance strategies but also for increase fundamental knowledge in the pathogenesis of phytoplasmas.

In Argentina, and other countries from South America, ‘*Candidatus* Phytoplasma meliae’ (16SrXIII-G, 16SrXIII-C) (Fernández et al. 2016) and ‘*Candidatus* Phytoplasma pruni’ (16SrIII-B) (Galdeano et al. 2004) are the causative agents of chinaberry decline and chinaberry yellows diseases, respectively. Despite the wide distribution that presents the host plant *Melia azedarach* L. along the Argentine territory, the distribution of ‘*Ca*. Phytoplasma meliae’ is restricted to North East while ‘*Ca*. pruni’ is widely represented throughout territory (Arneodo et al. 2007; Fernandez 2015). One of the factors that we believe would be modulating this pattern is the distribution of their vectors. Identifying and characterizing the Amp protein in these phytoplasmas constitutes the first step to achieve a more precise detection of potential vectors. The production of a specific Amp antisera which could be used as a diagnostic tool to survey potential vectors and also to evaluate the role of Amp in the transmission processes.

## Acknowledgments

This work was founded by INTA, MinCyT (Foncyt 2010-0810 and 2016-0862). We gratefully acknowledge Humberto Debat (IPAVE-CIAP-INTA) for valuable discussion and critical review of the manuscript.

## Compliance with ethical standards

The authors bear all the ethical responsibilities for this manuscript.

## Human and animal rights

The authors declare that the presented research does not include any animal and/or human trials.

## Informed consent

All authors consent to this submission.

---

**Figure S1.**
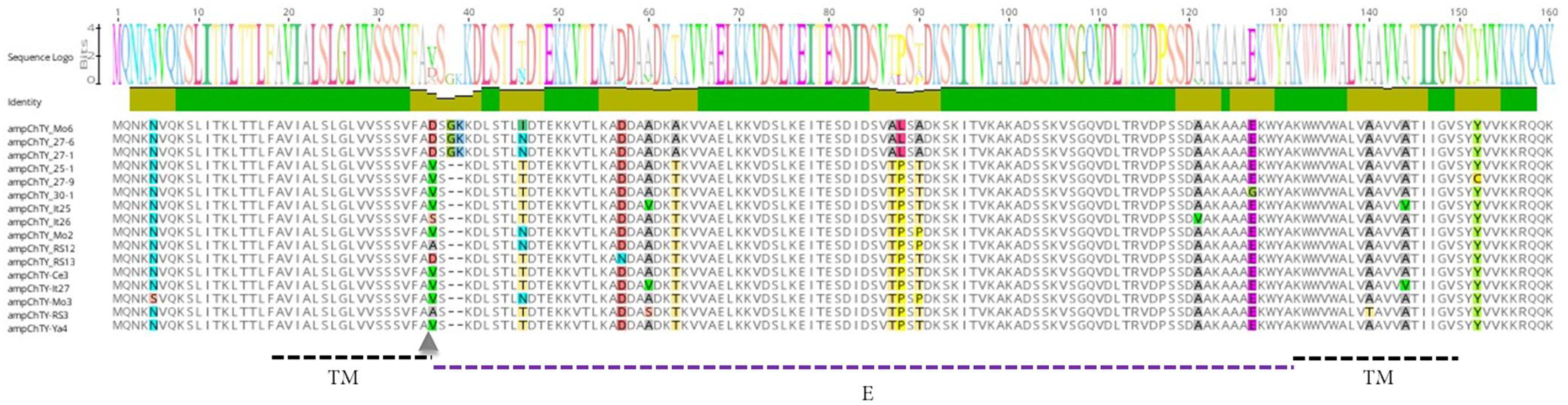
Multiple alignments of Amp amino acid residues in all ‘*Ca*. Phytoplasma meliae’ isolates obtained in this work. The TM domains are marked in black dotted lines while extracellular domain is marked in purple dotted lines, grey triangle shows the putative cleavage site

